# Boldness and exploration vary between shell morphs but not environmental contexts in the snail *Cepaea nemoralis*

**DOI:** 10.1101/866947

**Authors:** Maxime Dahirel, Valentin Gaudu, Armelle Ansart

**Author notes:** **Data accessibility** Data and code to reproduce all analyses are available on GitHub (https://github.com/mdahirel/cepaea-personality-2017) and archived in Zenodo (DOI: 10.5281/zenodo.3899042; version 1.1).

## Abstract

Understanding the maintenance of among-individual behavioral variation in populations, and predicting its consequences, are key challenges in behavioral ecology. Studying the association between repeatable behaviors and other traits under selection may shed light on the underlying selective pressures. We used the model snail *Cepaea nemoralis* to examine whether individual behavior is associated with shell morph, a key trait that has been extensively studied in the context of thermal tolerance and predator avoidance, and which is known to be under strict genetic control in this species. We quantified proxies of boldness and exploration in snails of three morphs coming from two habitats with different thermal contexts. We show that both behaviors were repeatable at the among-individual level (within-state *R*_*boldness*_ = 0.22 [95% credible interval: 0.15, 0.29]; *R*_*exploration*_ = 0.20 [0.15, 0.25]). Behavior was associated with shell morph, with the darker morph (five-banded) being consistently shyer and slower to explore. There was no evidence that thermal environment of origin influenced behavior. Snails became faster when test temperature increased; we found no evidence morphs differed in their thermal response. Boldness and exploration were correlated among individuals, forming a syndrome (*r* = 0.28 [0.10, 0.46]). We discuss what these results may tell us about the type of selection exerted by predators. We also detail how our results hint to a genetic link between shell morph and behavior, and the evolutionary implications of such a link. Finally, we discuss how our findings combined with decades of evolutionary research make *C. nemoralis* a very valuable model to study the evolution of behavior in response to environmental changes.

## Introduction

A key question in behavioral ecology, and more broadly in evolutionary ecology, is how to explain the persistence of variation in phenotypic traits. Although behavior is often seen as highly labile and dynamically adjustable to experienced conditions, individuals of many animal species exhibit “personalities”, i.e. behave consistently across time and contexts, and differ consistently from each other (Kralj-Fišer & Schuett, 2014; Réale, Reader, Sol, McDougall, & Dingemanse, 2007; Sih, Bell, Johnson, & Ziemba, 2004). This among-individual variation persists even when better adjustments of behaviors to environmental conditions would be adaptive, and the ability to tune behavior to conditions may itself vary among individuals (variation in “behavioral reaction norms”; Dingemanse, Kazem, Réale, & Wright, 2010). Moreover, behaviors are often correlated with each other and with other traits, forming multivariate syndromes (Réale et al., 2007; Royauté, Berdal, Garrison, & Dochtermann, 2018; Santostefano, Wilson, Niemelä, & Dingemanse, 2017), further constraining the range of behavioral phenotypes that are on display in populations (Dochtermann & Dingemanse, 2013).

State-dependent behavior is often invoked as one of the key mechanisms/frameworks potentially explaining both adaptive correlations/feedbacks between behaviors and other traits, and the maintenance of among-individual variation (Sih et al., 2015; Wolf & McNamara, 2012; Wolf & Weissing, 2010). Individuals can differ in morphology, size, past experienced environment, or any other so-called “state variables”, typically less labile than behavior or even fixed at the individual level. If the costs and benefits of behaviors vary depending on these state variables, then we should expect individuals differing in state to adaptively differ in behaviors as well (Wolf & Weissing, 2010). The pace-of-life hypothesis, which ties several axes of behavioral variation to underlying differences in life history and metabolism along a fast-slow axis (Réale et al., 2010; Wolf & McNamara, 2012; Wright, Bolstad, Araya-Ajoy, & Dingemanse, 2019), can be seen under this lens. Other examples include cases of phenotypic compensation, where predation risk can either be mitigated by behavioral changes or morphological defenses, leading to a positive association between risk-taking behavior and defenses (e.g. Ahlgren, Chapman, Nilsson, & Brönmark, 2015; but see De Winter, Ramalho Martins, Trovo, & Chapman, 2016 for a contradictory example). In some cases, quantitative genetics and/or experimental evolution approaches may provide evidence of the evolution of state-behavior associations (e.g. Kern, Robinson, Gass, Godwin, & Langerhans, 2016). In other cases in which this may be difficult, we believe that studying the association between personality and state traits can still provide valuable insights, especially if (i) the state trait is known to be fully genetically determined with little to no plasticity, (ii) we are able to study behavioral variation across a range of environments known to select on the state variable.

The grove snail *Cepaea nemoralis* (Linnaeus 1758) (family Helicidae) is a simultaneous hermaphrodite, medium-sized land gastropod common in western Europe (adult shell diameter 18-25mm; Welter-Schultes, 2012). It has a long history as a model in evolutionary biology, due to its conspicuous shell polymorphism (reviewed by Jones, Leith, & Rawlings, 1977; Ożgo, 2009)(Fig. 1A-B). Genetic variation in shell background color (from pale yellow to brown, but usually divided in yellow, pink, and brown; Davison, Jackson, Murphy, & Reader, 2019) and in the number or width of dark bands has been well described (Jones et al., 1977). Shell polymorphism is governed by a limited number of loci with a limited number of alleles (Richards et al., 2013), and by all evidence shows no phenotypic plasticity. Modern genomics studies now aim to pinpoint the actual molecular/physiological underpinnings of shell color (Kerkvliet, Boer, Schilthuizen, & Kraaijeveld, 2017; Richards et al., 2013).

**Figure 1.**
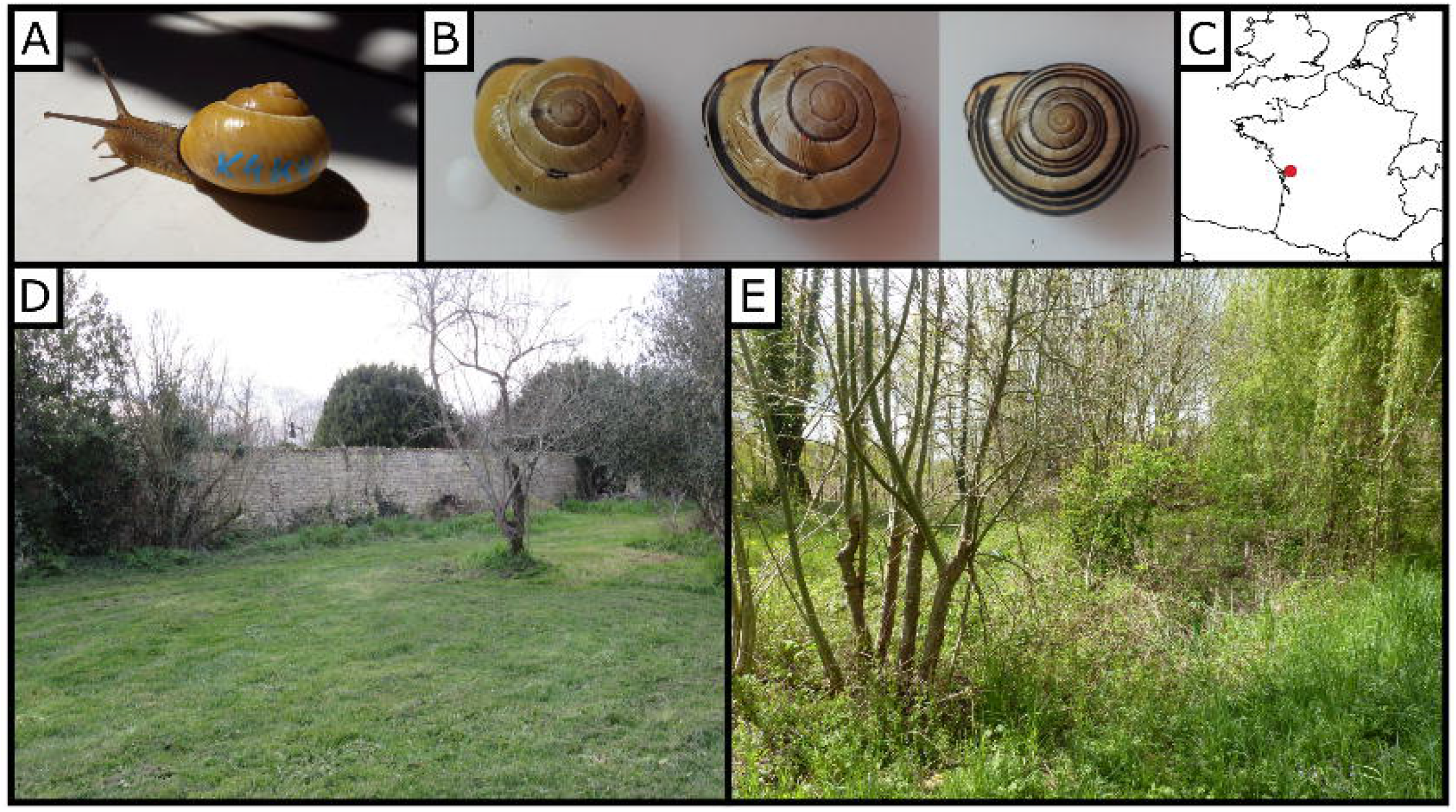
Study species and sites. (A) An unbanded yellow *Cepaea nemoralis*, showing the position of individual paint marks on the shell (B) Representative shells of the three studied morphs as seen from above: yellow unbanded, three-banded, and five-banded snails (C) Study sites location in France; the open habitat (D) and the shaded habitat (E) are separated by about 2 km. Photographs were taken during winter, when snails were collected.

In *C. nemoralis*, lighter-colored shells absorb less heat and allow snails to maintain a lower body temperature (Heath, 1975) and higher water content (Chang, 1991). Many studies have shown that lighter (vs. darker) snails have a selective advantage in hotter/sunnier (vs. colder/shaded) environments, whether one looks at continental-scale latitudinal clines (Jones et al., 1977; Silvertown et al., 2011), local-scale habitat comparisons (Kerstes, Breeschoten, Kalkman, & Schilthuizen, 2019; Ozgo & Kinnison, 2008; Schilthuizen, 2013), or historical comparisons in the context of climate change (Ożgo, Liew, Webster, & Schilthuizen, 2017; Ożgo & Schilthuizen, 2012). Local variations in morph frequencies have also been linked to predation pressure, generally in the context of visual selection (frequency-dependent selection and/or crypsis; Jones et al., 1977; Surmacki, Ożarowska-Nowicka, & Rosin, 2013, and references therein; but see Cook, 2008), but morph differences in shell resistance to crushing have also been described (Rosin, Kobak, Lesicki, & Tryjanowski, 2013).

Although this has been much less studied, shell morphs also vary in their behavior (Chang, 1991; Jones, 1982; Ożgo & Kubea, 2005; Rosin et al., 2018). Part of this variation is likely the direct consequence of differences in shell thermal properties and thus effective body temperature, as demonstrated by experiments that created “artificial” morphs by painting shells (Tilling, 1983). However, they probably also reflect, at least partly, intrinsic physiological differences: preferred temperatures can be altered using opioid agonists or antagonists, but banded snails are less responsive to this pharmaceutical manipulation (Kavaliers, 1992). Existing studies, however, have several major shortfalls for our understanding of the association between *Cepaea* morphology and behavioral syndromes. In particular, individuals are generally assayed once, which means separating within-from among-individual variation is impossible (Dingemanse & Wright, 2020; Niemelä & Dingemanse, 2018). This also means the level of total among-individual variation, and how it compares with among-morph variation, has remained to our knowledge unstudied. Additionally, all snails are often sampled from the same habitat, or habitat information is not used in behavioral analyses, meaning there is often no way to determine how behavior responds to selection pressures on shell color.

In this context, we investigated the existence and magnitude of personality variation and behavioral syndromes in *C. nemoralis*, how behavior is linked to shell variation, and how it is influenced by the environment of origin (sun-exposed or shaded) and currently experienced conditions. More specifically, we make the following hypotheses:

i. Exploration and boldness (risk-taking behavior) are both repeatable in this species, and positively correlated in a behavioral syndrome (Réale et al., 2010).
ii. As boldness may increase predation risk (e.g. Hulthén et al., 2017), we may expect phenotypic compensation through shell characteristics to be present in bolder individuals. This would lead to correlations between morph and behavior, the direction of which providing insights into the dominant selection pressures.
iii. As snails are ectotherms, exploration should increase with temperature due to increased metabolism (over the range of temperatures suitable to movement; Abram, Boivin, Moiroux, & Brodeur, 2017; Cloyed, Dell, Hayes, Kordas, & O’Gorman, 2019). We expect this temperature-exploration reaction norm should vary both in its slope and average value according to shell morph and habitat of origin. Populations having evolved in sun-exposed habitats, and lighter (unbanded) snails should be better adapted to maintain activity in the face of high temperatures (e.g. Cloyed et al., 2019), at the possible costs of lower activity at lower temperatures (Tilling, 1983).

## Methods

### Sampled sites and snail maintenance under laboratory conditions

Snails were sampled in fall 2016 in and close to the village of Arçais, France (Fig. 1C), roughly in the middle of the recorded latitudinal range of *Cepaea nemoralis* (GBIF Secretariat, 2020). We studied two sites located about 2 km apart and differing in terms of vegetation cover. One was a garden with few isolated trees, and thus under relatively direct sun exposure all year long (hereafter the “open habitat”; approximate location: 46° 17’ 50”N, 0° 41’ 30” W, Fig. 1D). The other was a 200 by 150 m deciduous forested lot, and thus fully shaded a large part of the year, especially the hottest spring and summer months (“shaded habitat”; approximately 46° 18’ 01” N, 0° 42’ 56” W, Fig. 1E). Only adult snails were selected (recognizable by a reflected “lip” on their shell opening), as a way to partly control for age. We only sampled snails with the three most abundant shell banding patterns: shells with no bands, three bands on the lower side of the shell, and five bands (Fig. 1B). Following previous authors (e.g. Kavaliers, 1992), we here focused for simplicity solely on band presence, and thus sampled only snails with yellow background shells, which are the most common in the study region (Silvertown et al., 2011; personal observations) and on which the contrast between shell background and dark bands is the strongest. We acknowledge that this may prevent us from fully generalizing, for now, to natural populations, as the effect of darker background color is not always the same as the effect of increased shell banding (e.g. Kerstes et al., 2019). Snails were hand-collected during the day, their period of inactivity, both by simplicity and to avoid skewing our sample towards more active individuals. If there were nonetheless a bias towards catching more conspicuous/ less likely to hide snails, we believe it would have artificially reduced, rather than increased, our effect sizes: we would have sampled the most active morph and the most active individuals from the least active morph, reducing mean morph differences.

Sampling for the present study was targeted and adjusted in the field to obtain roughly equal numbers of each banding pattern from both landscapes; it therefore did not allow us to make inferences on their relative abundances. The same sites were however sampled again in 2018 for a separate experiment, this time with random sampling relative to banding pattern. As in previous studies (e.g. Schilthuizen, 2013) and reflecting potential thermal selection, the darker five-banded snails were more frequent in the shaded habitat than in the open habitat (22.3 vs. 13.5 %; see Supplementary Material S1).

We transferred snails to the lab and kept them under dormancy conditions (6 ± 1°C, no light, food or water sources) until March 2017, about 3 weeks before the start of the experiment. We then divided them into groups of 15 individuals from the same landscape, five (randomly chosen) of each shell phenotype. Comparing group size to natural densities is difficult, due to the way natural densities are often reported in the literature (averages over entire habitats, including empty areas). However, groups of 10-20 individuals are commonly seen in the wild (personal observations) and are also often used in experiments (Oosterhoff, 1977; Rosin et al., 2018; Wolda, 1967). Groups were kept under controlled conditions (20 ± 1°C, L:D 16:8) in 8.5 × 15 × 12 cm polyethylene boxes lined with 1-2 cm of soil kept humid at the bottom. Snails had *ad libitum* access to prepared snail food (cereal flour supplemented with calcium, Hélinove, Saint Paul en Pareds, France) in a Petri dish. We gave each snail a unique ID written on the side of their shell with a paint marker (uni Posca, Mitsubishi Pencil Co., Ltd, Tokyo, Japan; Henry & Jarne, 2007). A total of 360 snails (60 for each habitat × shell phenotype combination) were used in the experiments described below. By necessity, the observer (see below) was not blind to individual habitat of origin/phenotype; note that the analyst (MD) did not contribute to the actual observations.

### Behavioral tests: boldness

We studied boldness using simulated predator attacks as in Dahirel et al. (2017). All tests were done by the same operator (VG) to avoid effects of inter-experimenter variability. Snails were assayed individually during the last four hours of the photophase, i.e. the early part of the daily activity period. Like other helicids, *Cepaea nemoralis* is nocturnal but tends to start activity sometime before dark (Cameron, 1970). To stimulate activity, we first placed them in a Petri dish with water for 5 minutes, before putting them on individual clean glass plates. After snails had moved at least one shell length (≈20 mm) from their starting position, the operator used a pipette tip to pinch them for 5 seconds on the right side of the foot. Preliminary tests confirmed that this was the shortest time needed to ensure all snails retracted fully in their shell. We then recorded the time snails took to exit the shell and resume activity after the attack (from retraction to the full extension of all tentacles out of the shell), as our measure of boldness (snails with shorter latencies being considered bolder). We stopped observations after 20 min if snails did not exit the shell. Snails from the same test box were tested on the same day, and placed back in their box after testing. To estimate the repeatability of boldness, snails were tested a second time after seven days, using the same protocol. The initial order in which groups were tested within a sequence was random; this order was conserved for all subsequent tests.

### Behavioral tests: exploration/speed

We studied snail movement at four temperatures within the activity range of *C. nemoralis* (Cameron, 1970): 15, 18, 22, and 25 °C. All tests were again performed by the same operator (VG), and again during the last four hours of the photophase each day. Movement tests started 7 days after the last boldness test for a given individual, successive movement tests were separated by 24h. Half of the boxes, equally distributed between landscapes of origin, were tested in increasing temperature order (from 15 to 25 °C), the other half in decreasing order (25°C to 15°). Twenty-four hours before a given test, we placed snails and their rearing box at the testing temperature for habituation, using temperature-controlled cabinets (ET 619-4, Lovibond, Dortmund, Germany). For testing, each snail was placed individually at the center of a clean 25 × 25 cm polyethylene box (height: 9 cm) and left free to move. Snails were deemed active once they had moved more than 2 cm away from their starting point. We used the time snails took to move more than 10 cm from their starting point, minus the time taken to start activity, as our exploration metric (with lower values for snails that moved away faster). We stopped observations after 20 min post-activity initiation. This metric was chosen for its ease of implementation; we acknowledge that it conflates exploration of the environment with movement speed (as both slow-moving individuals and thorough explorers would have higher first-passage times).

### Ethical note and compatibility with reporting guidelines

This study complies with all relevant national and international laws, and the ASAB/ABS Guidelines for the use of animals (2020) were adhered to as closely as possible. Potentially stressful experimental treatments (boldness experiment) were limited to the shortest possible time to elicit the behaviors of interest. No ethical board recommendation or administrative authorization was needed to work on or sample *Cepaea nemoralis*. The marking method used is non-invasive and has minimal to no documented effects on life-history traits (Henry & Jarne, 2007). We do not believe there is any potential for bias due to social background, self-selection, experience or other factors indicated in the STRANGE framework (Webster & Rutz, 2020). To the best of our knowledge, the studied individuals are representative of the local populations studied, except for the two constraints explicitly imposed on collection by our experimental design (only adults, equal numbers of a few morphs of interest). All individuals were subjected to the same experimental conditions once collected.

### Statistical analyses

We analyzed snail behavioral data in a Bayesian multilevel/mixed model framework, using the Stan language (Carpenter et al., 2017), with R (version 4.0; R Core Team, 2020) and the *brms* R package (Bürkner, 2017) as frontends. Scripting, analysis, and plotting relied on the *tidybayes*, *bayesplot*, and *patchwork* packages, as well as the *tidyverse* family of packages (Gabry, Simpson, Vehtari, Betancourt, & Gelman, 2019; Kay, 2019; Pedersen, 2019; Wickham et al., 2019).

We used a bivariate generalized linear multilevel model to estimate the effect of shell phenotype, habitat and temperature on behavior, quantify behavioral (co)variances and partition them across hierarchical levels (among-box, among-individual and within-individual variation) (Dingemanse & Dochtermann, 2013; Houslay & Wilson, 2017). We did not estimate within-individual trait correlations, as exploration and boldness were tested independently at the within-individual level (that is, boldness measure 1 had no stronger “link” to exploration measure 1 than boldness measure 2; scenario 4 of table 2 in Dingemanse & Dochtermann, 2013). We present a full write-up of the model as Supplementary Material S2; a general description follows below.

Boldness and exploration were analyzed assuming a lognormal distribution to account for the skewed distribution of time to event data. We accounted for the fact that monitoring was stopped before some individuals could express the behavior of interest by including a censored data indicator in the model. Fixed effects for both behaviors included shell banding (three-level categorical variable), landscape of origin (binary variable), and their interaction, as well as test order (1 or 2 for boldness, 1 to 4 for exploration). The model for exploration additionally included a test temperature effect as well as its interactions with shell banding and landscape. Categorical variables (shell banding, landscape of origin) were converted to centered dummy variables, and numeric variables (test order, temperature) were centered, following Schielzeth (2010)(temperature was additionally scaled to unit 1SD). This has two benefits. First, it makes main effect coefficients directly interpretable even in the presence of interactions (Schielzeth, 2010). Second, for categorical variables, having the intercept on an “average” rather than on one arbitrary default category avoids the problem of putting a more precise prior on an arbitrary reference category (which would be defined by the intercept only) than on the others (which would be defined by the intercept and one or several other coefficients)(McElreath, 2020). Morph-specific coefficients (intercepts, slopes) remain easy to obtain post-fitting, by simply adding the relevant posterior coefficients. Random effects included box-level and individual-level intercepts as well as, in the case of exploration, the associated slopes for temperature. This allowed us to estimate among-box and among-individual variation in mean behavior and thermal behavioral reaction norms as well as the box- and individual-level covariances among them (Dingemanse & Dochtermann, 2013).

We used a Normal(μ = ln(400), σ = 0.5) prior for the fixed effects intercepts (mean log-latencies), so that ~99% of the probability mass was within the range of latencies that was observable during the experiment (i.e. 0 to 1200 sec, see above), but not excluding larger values, because of censoring. We set the other priors to be weakly informative and follow some suggestions by McElreath (2020): a Normal(0,1) prior for the other fixed effects, a half-Normal(0, 1) prior for both random effect and distributional standard deviations. For the random effects correlation matrices, we use an LKJ(η = 3) prior, as it helps reach convergence faster than McElreath (2020)’s η = 2 default. Note that our choice here is more skeptical of high correlations and thus penalizes against our hypotheses of interest (there are detectable correlations).

We partitioned total phenotypic variation *V*_*P*_ for each behavior into the following components: *V*_*P*_ = *V*_*F*_ + *V*_*I*_ + *V*_*B*_ + *V*_*D*_, where *V*_*F*_ is the fixed effect variation, including *V*_*F(state)*_ the portion of fixed-effect variance attributable to known individual state (banding pattern, environment of origin), i.e. excluding experimental effects (test order, temperature) (estimated following de Villemereuil, Morrissey, Nakagawa, & Schielzeth, 2018); *V*_*I*_ the average among-individual variation (including the effect of random temperature slope, estimated following Johnson, 2014), with *V*_*I(intercept)*_ the among-individual variation at the average test temperature (*V*_*I*_ = *V*_*I(intercept)*_ for boldness); *V*_*B*_ and *V*_*B(intercept)*_ are the equivalent box-level variances; and *V*_*D*_ is the distributional, or residual, variation. As pointed by Wilson (2018) and de Villemereuil et al. (2018), there is in most cases no one “true” repeatability estimate just as there is no one “true” way of partitioning the phenotypic variance pie; several estimates with differing interpretations can be presented. Therefore, both absolute variance components and analytical choices regarding repeatabilities should be made explicit. We estimated the following two unadjusted repeatabilities (i.e. including the entirety of *V*_*P*_ in the denominator; Nakagawa & Schielzeth, 2010): within-state repeatability *R*_*(within-state)*_ = *V*_*I(intercept)*_ /*V*_*P*_, and what we term total repeatability, *R*_*(total)*_ = (*V*_*I(intercept)*_ + *V*_*F(state)*_)/*V*_*P*_. The proportion of persistent among-individual variation that is attributable to individual state (banding and landscape of origin) is then denoted by *V*_*F(state)*_ / ( *V*_*I(intercept)*_ + *V*_*F(state)*_). Variance components and repeatabilities are presented on the observed data scale (sensu de Villemereuil, Schielzeth, Nakagawa, & Morrissey, 2016). Variance components on the latent log scale (i.e. directly using model coefficients) led to qualitatively and quantitatively similar results.

We ran four chains for 12000 iterations, with the first 2000 iterations of each chain used for warmup. We checked mixing graphically and confirmed chain convergence using the improved 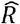 statistic by Vehtari et al. (2020). The chains were run longer than the default number of iterations to ensure the effective sample size was satisfactory for all parameters (both bulk- and tail-effective sample sizes sensu Vehtari et al., 2020 at least > 400, here > 1000). All posterior summaries are given as mean [95% highest posterior density interval].

## Results

Exploration was related to shell morph (Table 1, Fig. 2), with morph-specific intercepts, i.e. mean log-latencies, for 0, 3 and 5-banded snails of 6.65 [6.59, 6.69], 6.64 [6.59, 6.69] and 6.71 [6.66, 6.76], respectively. Five-banded snails were on average slower to explore their surroundings than either three-banded or unbanded snails (in both cases, mean difference: 0.07 [0.01, 0.13])(Fig.2) Snails also became slower as tests went on (Table 1). We found no clear evidence of an effect of the landscape of origin on exploration. Snails explored faster with increasing temperature (Table 1, Fig. 2; temperature slopes for 0, 3, and 5-banded snails: −0.12 [−0.16,−0.07], −0.11 [−0.16, −0.07], −0.10 [−0.15,−0.06]). There was however no clear evidence that the slope of the temperature reaction norm varied between the three morphs, or between snails coming from different landscapes (Table 1; credible intervals for all interactions largely overlap 0).

**Table 1:**
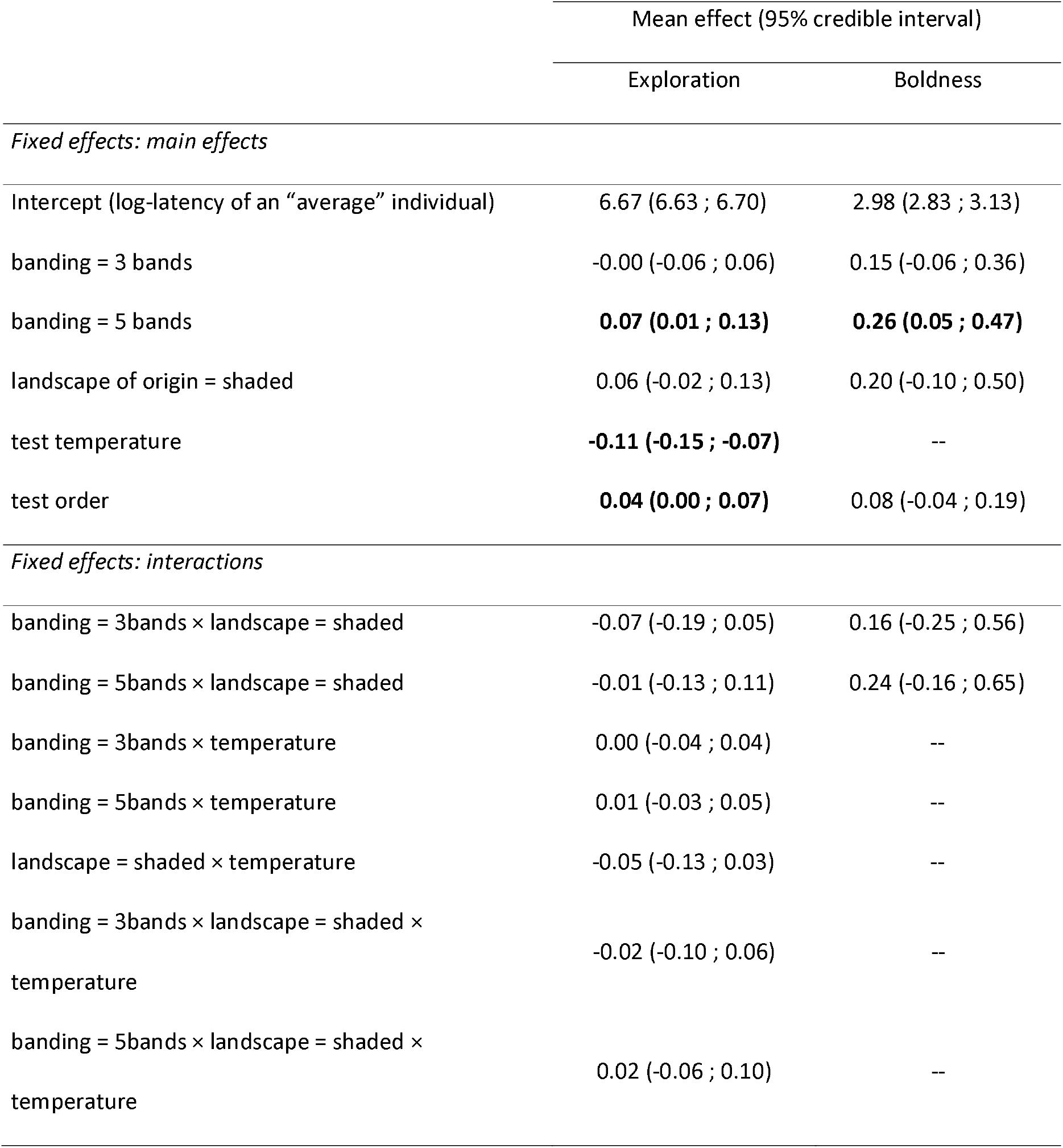
Estimated fixed effect parameters of the model explaining exploration and boldness latencies (mean and 95% credible intervals). Continuous explanatory variables are centered and scaled, and categorical variables converted to centered dummy variables; the intercept then refers to the behavior of a hypothetical “average” snail.

**Figure 2.**
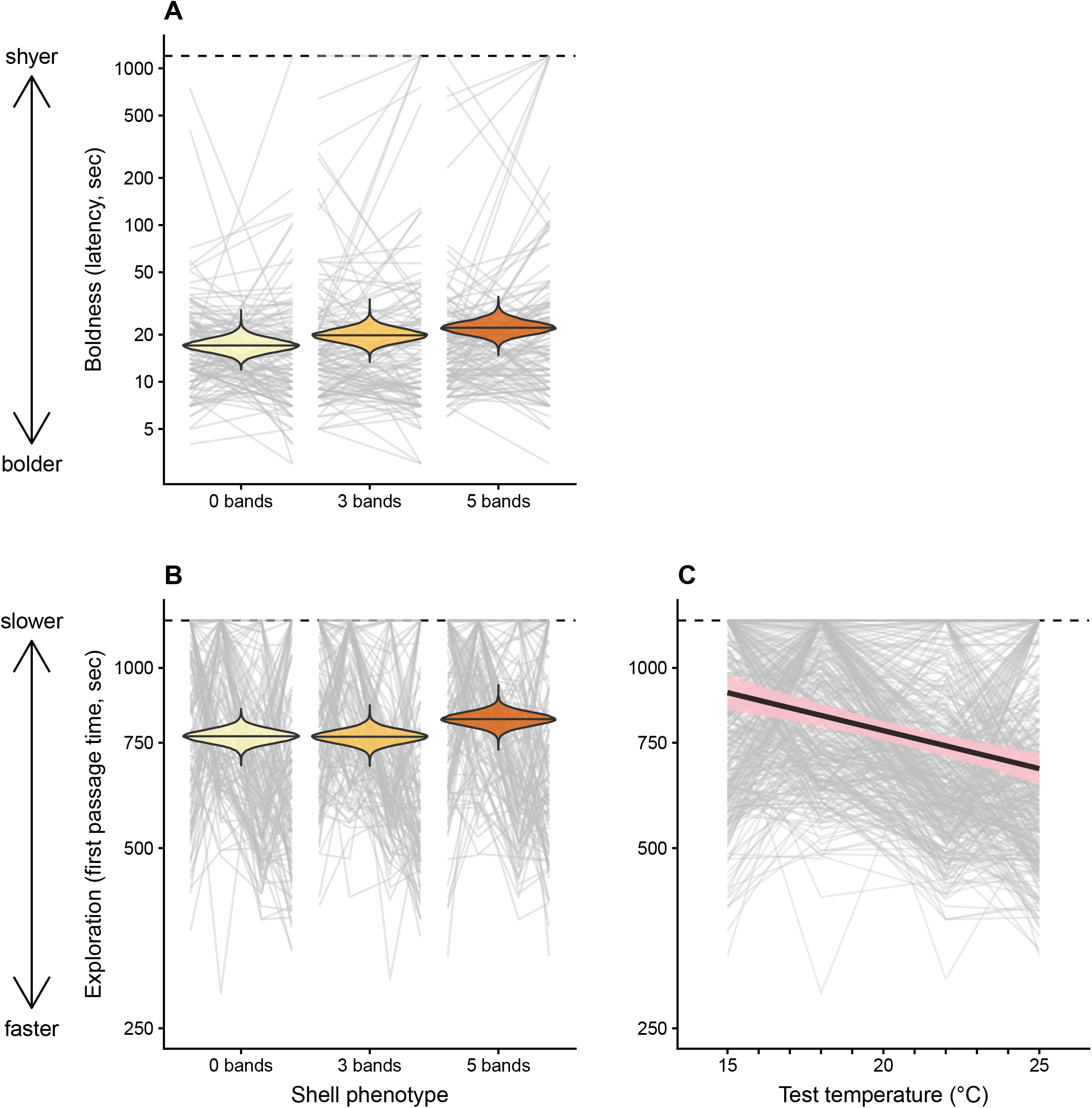
(A; B) posterior distributions of median boldness (A) and exploration (B) in relation to shell morph. The model estimates the mean log-latencies, which correspond to the medians on the observed latencies scale. (C) Mean and 95% credible band for the relationship between exploration latency and test temperature. Grey lines connect trials from the same individual. Values are plotted on a log-transformed axis. N = 360 individuals (60 per landscape × morph combination)

Morphs also varied in average boldness (Table 1, Fig. 2), with morph-specific intercepts for 0, 3 and 5-banded snails of 2.84 [2.65, 3.04], 2.99 [2.79, 3.18] and 3.10 [2.90, 3.30], respectively. Unbanded snails were bolder than five-banded snails (mean difference: −0.26 [−0.47, −0.05]); three-banded snails presenting intermediate values, with no clear difference with either extreme morph. Again, there was no evidence for landscape or landscape × morph effects.

Both exploration and boldness were repeatable at the individual level, with average repeatabilities in the same range for both behaviors (Table 2). Including fixed effect variation due to individual state (morph and landscape of origin) in the calculation only slightly increased repeatabilities. Indeed, the proportion of persistent among-individual variation attributable to fixed effects was different from zero but small, with over 90% of individual-level variation attributable to other, unmeasured sources (Fig. 3, Table 2). Among-individual variation in temperature slopes was minimal, with variation in intercepts explaining 98% [89%, 100%] of the average exploration *V*_*I*_ (Fig. 3, Table 2). Accordingly, we find no clear evidence that the level of among-individual variation changes across the temperature gradient; following equations in Brommer (2013), the ratio between latent-scale *V*_*I*_ at the lowest and highest tested temperatures is not different from 1 (0.76 [0.39, 1.13]). We also found no evidence of widespread rank switching across the temperature gradient (faster than average individuals in one environment remained overall faster across contexts): indeed, the cross-environmental correlation, which is higher the more predicted individual rankings stay consistent across environmental gradients (Brommer, 2013), was close to 1 when comparing the two extremes of the thermal gradient (0.85 [0.62, 1.00] on the latent scale).

**Figure 3.**
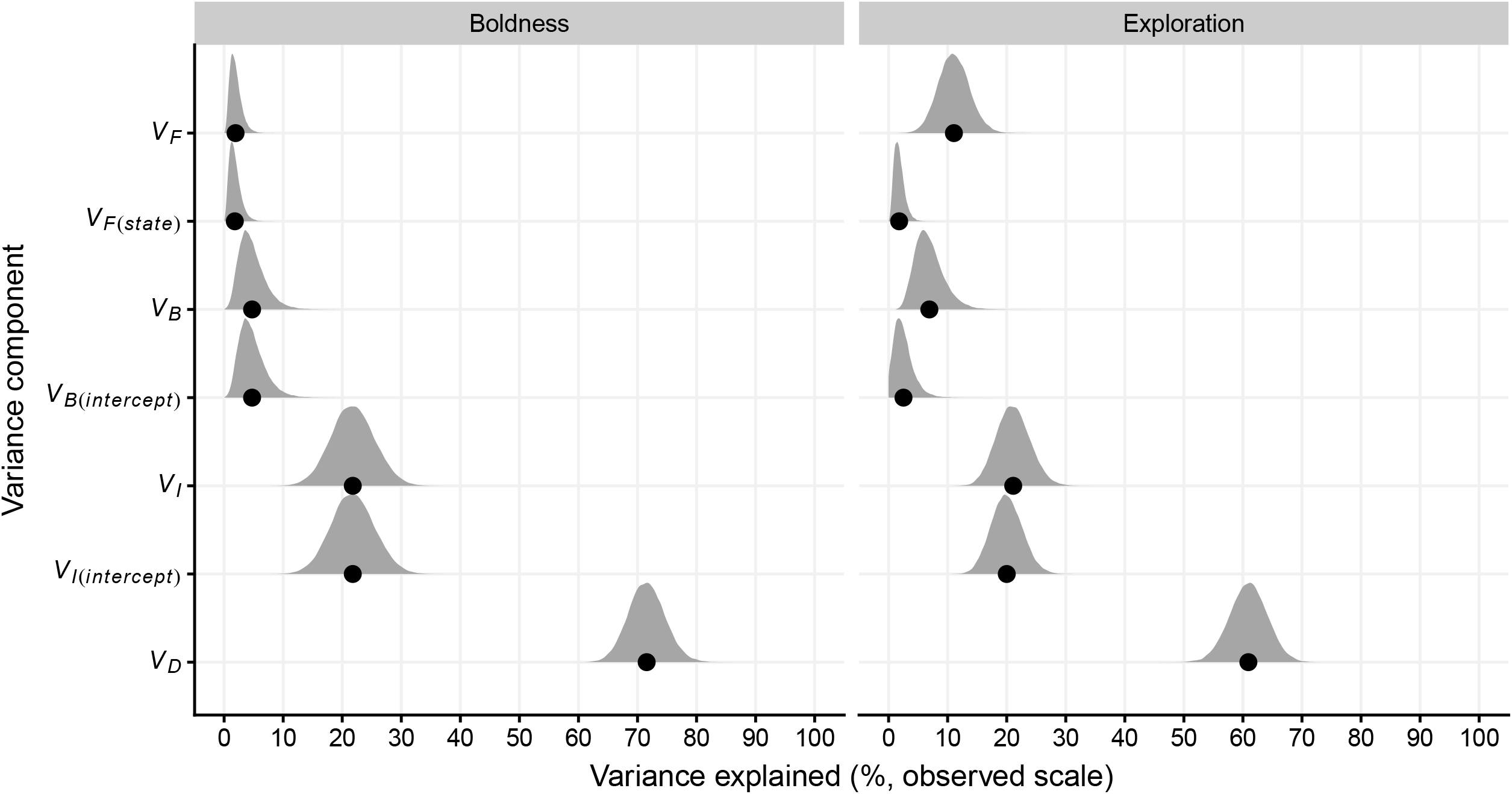
Mean (points) and posteriors for the proportion of variance explained by the different variance components. For boldness, *V*_*B*_ and *V*_*B(intercept)*_ are exactly equal by definition (same for *V*_*I*_ and *V*_*I(intercept)*_); see Methods and Table 2 for details on this and the names of the variance components.

**Table 2.**
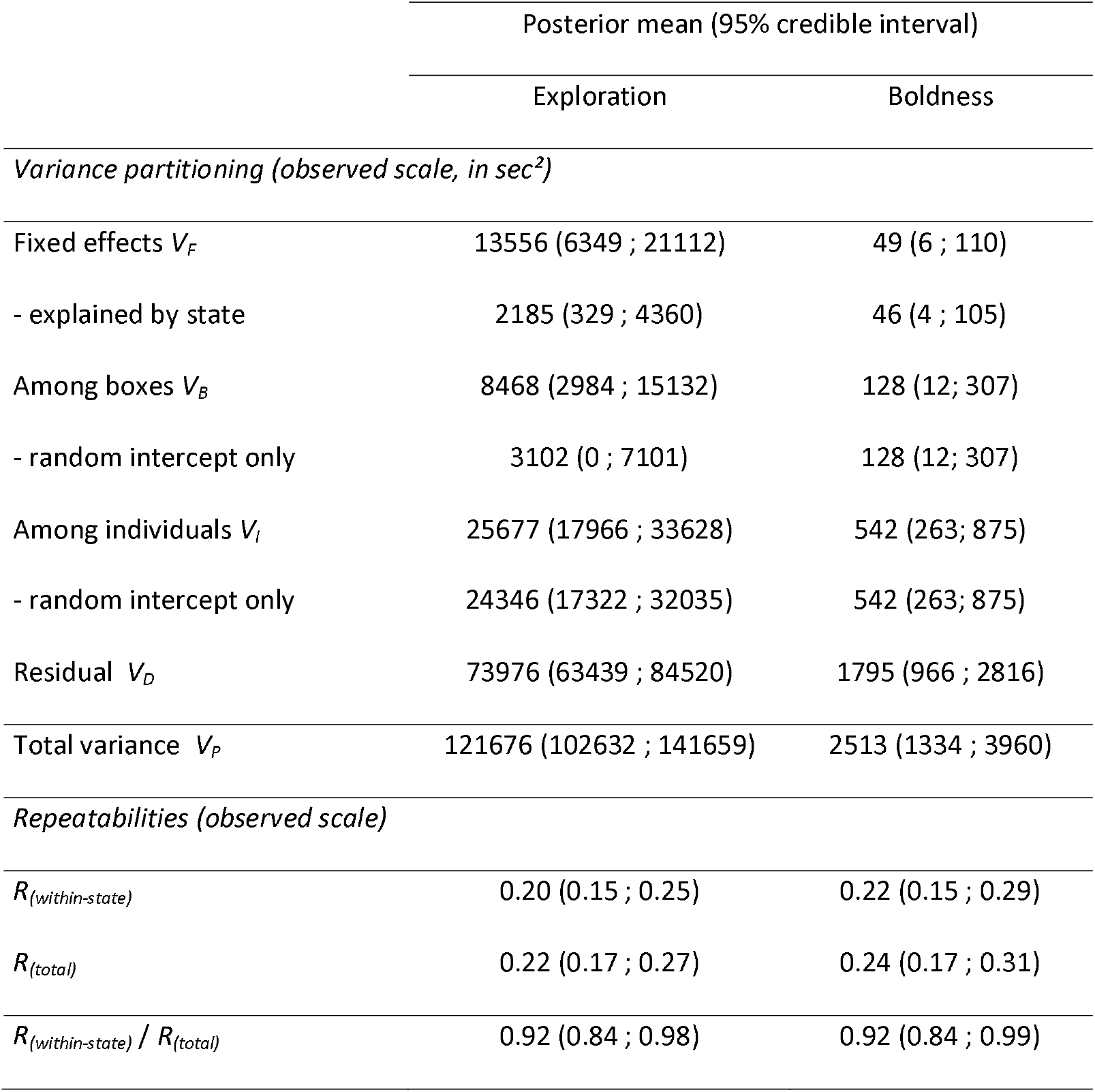
Variance partitioning and repeatabilities. Estimated variances and 95% credible interval are obtained from a multivariate mixed model. Variances are estimated on the observed scale *sensu* de Villemereuil et al (2016). Variances are rounded to the nearest unit. Readers looking at the data may note that the total variances *V*_*P*_ are higher than the empirically observed variances. This is because the latter are underestimated, due to censoring.

Variation among boxes was small but non-negligible, in the same range as the proportion of variation explained by fixed effects for both behaviors. Average exploration and boldness were positively correlated at the individual level (Table 3, Fig. 4). There was no evidence that among-individual variation in responses to temperature was correlated with either mean exploration times or mean boldness (Table 3). There was no evidence for box-level correlations among traits (Table 3).

**Table 3.** Random effect correlation matrices (latent scale, mean and 95% credible intervals). Box-level correlations are above the diagonal, individual-level correlations below the diagonal.

|  | Boldness, intercept | Exploration, intercept | Exploration, temperature slope |
| --- | --- | --- | --- |
| Boldness, intercept |  | 0.12 (-0.42 ; 0.63) | 0.38 (-0.05 ; 0.79) |
| Exploration, intercept | **0.28 (0.10 ; 0.46)** |  | -0.19 (-0.69 ; 0.33) |
| Exploration,<br>temperature slope | -0.19 (-0.67 ; 0.32) | 0.29 (-0.24 ; 0.76) |  |

**Figure 4.**
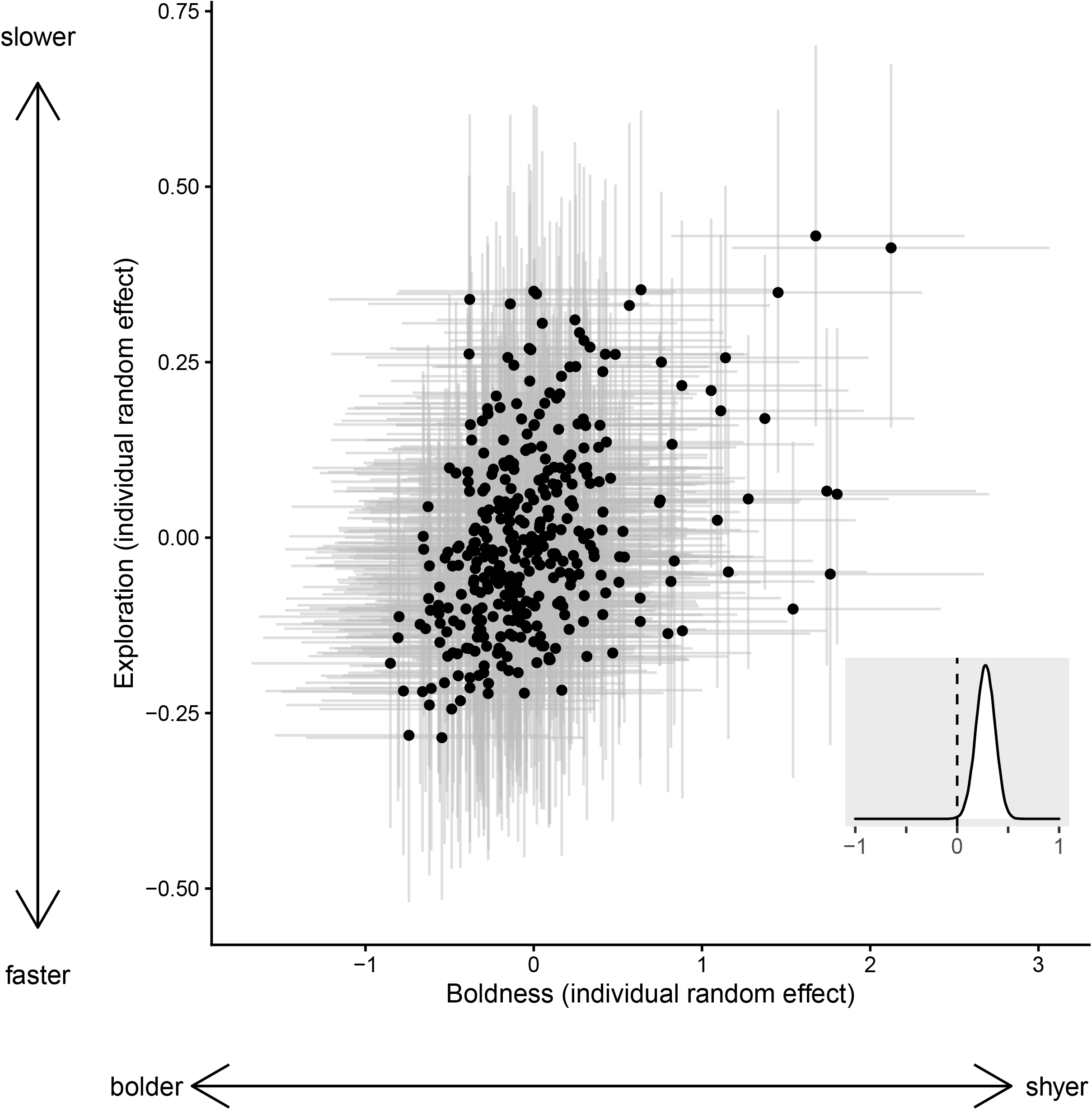
Correlation between individual-level random effects for boldness and exploration, illustrated by plotting their respective Best Linear Unbiased Predictors and 95% credible intervals. Inset: posterior distribution of the correlation coefficient. BLUPs are plotted, and correlation estimated, on the latent log scale.

## Discussion

By showing that behaviors linked to boldness and exploration are repeatable in *Cepaea nemoralis*, we add to a growing list of personality studies in gastropods, highlighting the usefulness of this taxon to address key questions in behavioral ecology (see e.g. Ahlgren et al., 2015; Cornwell, McCarthy, & Biro, 2020; Cornwell, McCarthy, Snyder, & Biro, 2019; Dahirel et al., 2017; Goodchild, Schmidt, & DuRant, 2020; Seaman & Briffa, 2015). We note however that this list is biased towards freshwater and marine gastropods; more studies are needed to understand among-individual variation in behavior in land mollusks. Additionally, we demonstrated that boldness and exploration are positively correlated in a common syndrome and that their expression varies depending on shell banding, a trait under strictly genetic determinism (little to no plasticity) that has been the focus of a lot of research in this species (Ożgo, 2009; Richards et al., 2013). Given how behavior can shape effective thermal tolerance (Abram et al., 2017) or vulnerability to predation (e.g. Hulthén et al., 2017), we believe these behavioral differences must be taken into account when discussing the evolution of shell color in this model species.

Unbanded snails were both bolder and explored faster than five-banded snails (Table 1, Fig. 2). Three-banded snails behaved similarly to unbanded snails for exploration (but were intermediate between unbanded and five-banded snails for boldness). This shows the “effectively unbanded” category sometimes used in *Cepaea* studies (Cain & Sheppard, 1954; Ożgo & Schilthuizen, 2012) has at least some behavioral relevance (that category groups together snails with little to no banding on the side of their shell exposed to the sun). Exploration and boldness were positively correlated both at the among-individual (Table 2) and among-morph levels (the shyest morph was also the slowest, Fig. 2). At the individual level, while some clutches were laid during the experiments, we were not able to test if this behavioral syndrome was integrated into a broader pace of life syndrome *sensu* Réale et al. (2010) by linking behavioral and life-history variation. Indeed, we were unable to ascertain the maternal and especially paternal origin of most clutches, and were not able to follow snail fecundity or longevity over their entire life. There are however some indications in the literature that more active/mobile snails are faster-growing (Oosterhoff, 1977), as the pace-of-life syndrome hypothesis would predict.

Five-banded snails were on average shyer than unbanded snails (Fig. 2). Birds, thrushes in particular (genus *Turdus*), are key predators of *Cepaea nemoralis* (Rosin, Lesicki, Kwieciński, Skórka, & Tryjanowski, 2017; Rosin, Olborska, Surmacki, & Tryjanowski, 2011). Historically, both frequency-dependent predation and direct visual selection due to crypsis have been invoked as explanations for predator-dependent morph variation in *Cepaea* (Jones et al., 1977; Ożgo, 2009), but discussions often used human vision as a baseline. More recently, crypsis explanations have received increased support from an experiment using models of avian vision to more rigorously test how thrushes see different shell morphs (Surmacki et al., 2013). In both our test sites, the boldest morph (unbanded shell) is the least conspicuous (based on Surmacki et al., 2013), not the rarest. Building on the phenotypic compensation hypothesis (i.e. that risk-taking individuals should be better defended; Ahlgren et al., 2015; Kuo, Irschick, & Lailvaux, 2015), this result then adds support to crypsis-based explanations of *Cepaea* morph variation. However, phenotypic compensation is not a hard rule, and risk-taking individuals are sometimes less defended than risk-avoiding ones (De Winter et al., 2016; Goodchild et al., 2020). Besides, snails are also predated by rodents (Rosin et al., 2011), and shell morphs differ in shell strength in ways that go counter to the phenotypic compensation hypothesis (5-banded shells being stronger; Rosin et al., 2013). The combined effect of color and shell thickness/strength on predation risk remains to be studied. Finally, we must remember that (i) our knowledge of how avian predators perceive snails is very limited (Surmacki et al., 2013), (ii) we only tested a small set of the available morphs, which do not include the rarest background colors (pink and brown), and (iii) shell banding is a trait under multiple selection pressures, including thermal selection (see below).

Exploration speed was temperature-dependent: as expected from an ectothermic species, snails were on average faster at higher temperatures (Fig. 2). The temperature reaction norm of exploration was remarkably conserved among individuals (the near-totality of the among-individual variance *V*_*I*_ was due to differences in average behavior, rather than in temperature slopes; Table 2, Fig. 3). In addition, there was surprisingly no evidence that behavioral differences among morphs are influenced by the thermal environment, whether we consider the environment of origin (no habitat × morph interaction) or the current environment (no effect of morph identity on thermal reaction norms) (Table 1). This is despite abundant evidence in the literature for thermal selection on shell morphs, based on both field comparisons (e.g. Richardson, 1974; Schilthuizen, 2013; Kerstes et al., 2019; for this study, see Methods), and experiments (Lamotte, 1959; Tilling, 1983; Wolda, 1967). Studies giving snails a choice between multiple temperatures show snail morphs do have different thermal preferences that align with expectations based on thermal selection (Kavaliers, 1992). Some studies suggest that snails use shade and humidity just as much (and potentially more) as temperature as cues to adjust their behavior to microclimate (Ożgo & Kubea, 2005; Rosin et al., 2018). Our exploration tests were short, under standardized lighting conditions and with no water, and snails were brought back to favorable humidity soon after. It is possible longer experiments, or experiments comparing the responses of snails from different habitats to realistic climate variation (including shade and/or humidity) would yield different responses. Maybe more importantly, we only tested temperatures favorable for activity, i.e. the limited part of the thermal niche closer to the optimum. Morph differences in behavior might be stronger closer to critical minimal or maximal temperature thresholds (Tilling, 1983). This can be investigated by using a wider range of temperatures and expanding the reaction norm approach used here to either a character state approach (e.g. Houslay, Earley, Young, & Wilson, 2019) or a non-linear reaction norm approach (Arnold, Kruuk, & Nicotra, 2019); both would account for the typical non-linearity of complete thermal performance curves (Arnold et al., 2019). It is very important to note, however, that these results do not mean populations from landscapes differing in sun exposure are identical in behavior, even for the range of situations we tested. Indeed, because morphs differ in their behavior, and because morph frequencies differ among landscapes (see Supplementary Material S1), the average snail from a sun-exposed population may well be bolder and more active than its counterpart from a shaded population.

In any case, the links between behaviors and morphs we observed are conserved across contexts, despite (apparent) selection on shell morph. While this is not a definite proof by itself, we consider this a first hint in favor of a genetic association between morphs and behaviors that cannot be easily broken by environmental changes. In addition to studies aiming to confirm these behavioral traits are heritable, further research into the physiological underpinnings of behavioral differences between morphs (building on e.g. Kavaliers, 1992) and of shell color and pattern determination (Kerkvliet et al., 2017) should help confirm (or infirm) this putative genetic correlation and elucidate its proximate basis.

Assuming this genetic link is confirmed, any discussion about how selection on morph may influence the evolution of behavior (or vice versa) must be tempered by one fact: the greater part of the repeatable among-individual variation in behavior was not explained by shell morph (see Fig. 3, Table 1, and the fact that morph differences are hard to see from raw data in Fig. 2). It is in a way unsurprising, as we did not expect a single discrete trait to entirely constrain individual behavioral variation. Indeed, the expression of animal personalities can be influenced by many unobserved drivers and state variables which should have a priori limited links to shell morph and its drivers (Burns et al., 2012; Petelle, Martin, & Blumstein, 2019; Sih et al., 2015; Wright et al., 2019). This includes for instance sex or reproductive history (DiRienzo & Aonuma, 2017; Kralj-Fišer, Hebets, & Kuntner, 2017), predation risk (Goodchild et al., 2020), age or life stage (Dahirel et al., 2017), or body size (Santostefano et al., 2017). Snail behavior is particularly sensitive to population density including during development (Cameron & Carter, 1979; Oosterhoff, 1977), an environmental axis we ignored in the present study. Also, our study focused on relatively short-term repeatability; it is possible that over larger time scales, the variance component related to morph differences plays a more important role. In a fish community, for instance, some differences among species are detectable over long but not short time scales (Harrison et al., 2019). Finally, some level of stochastic behavioral individuality is inevitable even in the total absence of meaningful genetic and environmental variation (Bierbach, Laskowski, & Wolf, 2017). What must be noted, though, is that some of this “remaining” among-individual variation may still, actually, relate to shell morph. Indeed, because of dominance within loci and especially epistatic relationships among loci, individuals that share the same shell phenotype may actually vary greatly in terms of underlying shell genotype (e.g. having one dominant allele at the « band presence » locus leads to total band absence and masks variation at all other banding genes; Jones et al., 1977). However, while much is known about among-morph variation in thermal tolerance, life history, physiology (Kavaliers, 1992; Kerstes et al., 2019; Lamotte, 1959; Oosterhoff, 1977; Richardson, 1974; Tilling, 1983; Wolda, 1967), we know nothing, as far as we can tell, about within-morph, but among-genotype variation. Investigations using individuals of known genotype obtained through repeated crosses or, as our knowledge of the actual molecular underpinnings increases, through direct genotyping (Gonzalez, Aramendia, & Davison, 2019; Kerkvliet et al., 2017), may shed light on this “hidden” genetic variation and whether it contributes to the persistence of morph-related behavior differences.

Increased boldness and exploration have been tied to a higher probability of dispersal in many species (Cote, Clobert, Brodin, Fogarty, & Sih, 2010), including land snails (Dahirel et al., 2017), and non-random dispersal is now acknowledged as a potentially widespread force behind population phenotypic divergence (Edelaar & Bolnick, 2012; Jacob, Bestion, Legrand, Clobert, & Cote, 2015). Bolder animals are often thought to trade increased success against a greater predation risk (Hulthén et al., 2017; but see Moiron, Laskowski, & Niemelä, 2020); predation is generally considered a key driver of morphological differences in *Cepaea*, and plays a key role in dispersal across taxa (Fronhofer et al., 2018). Although active dispersal can safely be dismissed as a driver of continental-scale differences in morph frequencies, our results point to *Cepaea* as a good model to understand how existing behavioral differences may drive local-scale morphological differences (and vice versa). By building on, and complementing, a decades-long history of evolutionary research, this will help us better understand the role of behavior, and constraints on behavioral variation, in shaping responses to rapid environmental changes (Candolin & Wong, 2012), including landscape alteration and climate change.

## Acknowledgments

We thank Youn Henry and Kévin Tougeron for helpful pointers on the link between behavior and thermal tolerance, as well as two anonymous referees and the editor for their comments on previous versions of this article.

