## Supplementary Material for "Boldness and exploration vary between shell morphs but not environmental contexts in the snail *Cepaea nemoralis*"

**Supplementary Material S1: Morph differences among the 2 studied landscapes**

Our sampling protocol for the experiment described in the main text did not allow us to estimate morph proportions. However, the same two landscapes were revisited in 2018, with this time a random sampling protocol with respect to banding pattern. We captured 336 snails overall (*N_tot_*_[shaded]_ = 166; *N_tot_*_[sun-exposed]_ = 170) and analysed the proportion of five-banded snails using a binomial generalized linear model:

$$N_{5b[\mathrm{landscape}]} \sim\text{Binomial}\left( n=N_{tot\left[ \mathrm{landscape} \right]}, p= p_{5b[\mathrm{landscape}]} \right)$$

(1)

$$logit\left( p_{5b[\mathrm{landscape}]} \right)= \beta_{0[landscape]}$$

(2)

, with $p_{5b[\mathrm{landscape}]}$ the expected proportion of five-banded snails in a given landscape. The model was fitted using Stan and *brms* (see main text) and a weakly informative prior for $\beta_{0[landscape]}$, Normal(μ= 0, σ = 1.5) (McElreath 2020).

We find that five-banded snails are more frequent in the shaded landscape (37/166, 22.3 %, posterior prediction: 22.5% [16.2%, 28.8%]) than in the open habitat (23/170, 13.5 %, posterior prediction: 13.9 [9.2%, 19.3%]), with a posterior predicted difference of 8.6% (95% credible interval: 0.1%, 16.9%).

**Supplementary Material S2: Formal description of the multivariate multilevel model**

The multivariate model used in the manuscript to analyse boldness and exploration in *Cepaea nemoralis* can be written formally as following (equations 1 to 16):

$$B_{ijk} | \left( C_{\left[ \mathrm{bold} \right]ijk}=0 \right) \sim\text{logNormal}\left( \mu_{[bold]ijk},\sigma_{0[bold]} \right)$$

(1)

$$B_{ijk} \left| {(C}_{\left[ \mathrm{bold} \right]ijk}=1 \right) \sim\text{logNormal}\text{-CCDF}\left( \mu_{[bold]ijk},\sigma_{0[bold]} \right)$$

(2)

$$E_{ijk}| \left( C_{\left[ \exp\right]ijk}=0 \right) \sim\text{logNormal}\left( \mu_{[exp]ijk},\sigma_{0[exp]} \right)$$

(3)

$$E_{ijk}\left| (C_{\left[ \exp\right]ijk}=1 \right) \sim\text{logNormal}\text{-CCDF}\left( \mu_{[exp]ijk},\sigma_{0[exp]} \right)$$

(4)

, where $B_{ijk}$ and $E_{ijk}$ are boldness and exploration latencies values of individual *i* in box *j* at trial *k*; the _[bold]_ and _[exp]_ subscripts denote whether a parameter corresponds to boldness or exploration, respectively; $C$ is a censorship indicator set to 1 if the behaviour was not observed by the end of the trial, 0 if it was; CCDF refers to the complementary cumulative density function used to estimate censored models (see e.g. Stan Development Team 2018); $\mu$ and $\sigma_{0}$ denote the means and standard deviations of the log-transformed latencies. Mean log-transformed latencies $\mu$ are assumed to depend on individual identity and experimental conditions as follow:

$$\mu_{[bold]ijk}=\beta_{0[bold]}+\sum_{h=1}^{p} \beta_{h[bold]}x_{hijk}+\alpha_{[bold]i} +\gamma_{[bold]j}$$

(5)

$$\mu_{[exp]ijk}=\beta_{0[exp]}+\sum_{h=1}^{q} \beta_{h[exp]}x_{hijk}+{\alpha_{\left[ \exp\right]i}+\kappa}_{[exp]i} t_{ijk}+\gamma_{[exp]j}+\lambda_{[exp]j}t_{ijk}$$

(6)

, where $\beta_{0}$ are the intercepts; $\beta_{h}$ the fixed effect coefficients associated with each explanatory variable $x_{h}$ (including interactions, see main text), $\alpha_{i}$ and $\gamma_{j}$ the individual- and box-specific average deviations from fixed effects, $\kappa_{i}$ and $\lambda_{j}$ the individual- and box-specific effects of temperature, and $t_{ijk}$ the (scaled) test temperature for the *k*^th^ exploration trial of individual *i* in box *j*. The random effects $\alpha$, $\gamma$, $\kappa$ and $\lambda$ are distributed according to the following multivariate normal distributions:

$$\left[ \begin{matrix} \gamma_{[bold]j} \\ \gamma_{[exp]j} \\ \lambda_{[exp]j} \end{matrix} \right] \sim MVNormal(\left[ \begin{matrix} 0 \\ 0 \\ 0 \end{matrix} \right] ,\boldsymbol{\Omega}_{\mathbf{box}})$$

(7)

$$\left[ \begin{matrix} \alpha_{[bold]i} \\ \alpha_{[exp]i} \\ \kappa_{[exp]i} \end{matrix} \right] \sim MVNormal(\left[ \begin{matrix} 0 \\ 0 \\ 0 \end{matrix} \right] ,\boldsymbol{\Omega}_{\mathbf{ind}})$$

(8)

$$\boldsymbol{\Omega}_{\mathbf{box}}= \left[ \begin{matrix} \sigma_{\gamma_{[bold]}} & 0 & 0 \\ 0 & \sigma_{\gamma_{[exp]}} & 0 \\ 0 & 0 & \sigma_{\lambda_{[exp]}} \end{matrix} \right] \mathbf{R}_{\boldsymbol{box}} \left[ \begin{matrix} \sigma_{\gamma_{[bold]}} & 0 & 0 \\ 0 & \sigma_{\gamma_{[exp]}} & 0 \\ 0 & 0 & \sigma_{\lambda_{[exp]}} \end{matrix} \right]$$

(9)

$$\boldsymbol{\Omega}_{\mathbf{ind}}= \left[ \begin{matrix} \sigma_{\alpha_{[bold]}} & 0 & 0 \\ 0 & \sigma_{\alpha_{[exp]}} & 0 \\ 0 & 0 & \sigma_{\kappa_{[exp]}} \end{matrix} \right] \mathbf{R}_{\mathbf{ind}} \left[ \begin{matrix} \sigma_{\alpha_{[bold]}} & 0 & 0 \\ 0 & \sigma_{\alpha_{[exp]}} & 0 \\ 0 & 0 & \sigma_{\kappa_{[exp]}} \end{matrix} \right]$$

(10)

$$\mathbf{R}_{\mathbf{box}}= \left[ \begin{matrix} 1 & r_{\gamma_{[bold]}\gamma_{[exp]}} & r_{\gamma_{[bold]}\lambda_{[exp]}} \\ r_{\gamma_{[bold]}\gamma_{[exp]}} & 1 & r_{\gamma_{[exp]}\lambda_{[exp]}} \\ r_{\gamma_{[bold]}\lambda_{[exp]}} & r_{\gamma_{[exp]}\lambda_{[exp]}} & 1 \end{matrix} \right]$$

(11)

$$\mathbf{R}_{\mathbf{ind}}= \left[ \begin{matrix} 1 & r_{\alpha_{[bold]}\alpha_{[exp]}} & r_{\alpha_{[bold]}\kappa_{[exp]}} \\ r_{\alpha_{[bold]}\alpha_{[exp]}} & 1 & r_{\alpha_{[exp]}\kappa_{[exp]}} \\ r_{\alpha_{[bold]}\kappa_{[exp]}} & r_{\alpha_{[exp]}\kappa_{[exp]}} & 1 \end{matrix} \right]$$

(12)

, where $\text{Ω}_{\text{box}}$ and $\boldsymbol{\Omega}_{\mathbf{ind}}$ are variance-covariance matrices for the box- and individual-level random effects, $\mathbf{R}_{\mathbf{box}}$ and $\mathbf{R}_{\mathbf{ind}}$ the corresponding correlation matrices, $\sigma$ denote random effect standard deviations and $r$ correlation coefficients.

Finally, we assume the following prior distributions (see main text for justifications):

$\beta_{0} \sim\text{Normal}(\ln\left( 400 \right),0.5)$

(13)

$\beta_{h}| (h>0) \sim\text{Normal}(0,1)$

(14)

$\sigma\sim\text{HalfNormal}(0,1)$

(15)

$\mathbf{R} \sim\text{LKJcorr}\left( \eta=3 \right)$

(16)

where priors for $\sigma$ and $\mathbf{R}$ refer to all standard deviations and correlation matrices, respectively.

Results are largely insensitive to the choice of prior parameters (not shown): for instance, changing the prior for $\beta_{0}$ to be identical to the one for the other $\beta$, and/or changing the standard deviation of the latter to up to 5 leads to the same results (within random fluctuations), and simply increases time to reach satisfactory effective sample sizes.

Stan Development Team 2018. Stan user guide, version 2.20.
